# A mosaic adeno-associated virus vector as a versatile tool that exhibits high levels of transgene expression and neuron specificity in primate brain

**DOI:** 10.1101/2021.07.18.452859

**Authors:** Kei Kimura, Yuji Nagai, Gaku Hatanaka, Yang Fang, Soshi Tanabe, Andi Zheng, Maki Fujiwara, Mayuko Nakano, Yukiko Hori, Ryosuke Takeuchi, Mikio Inagaki, Takafumi Minamimoto, Ichiro Fujita, Ken-ichi Inoue, Masahiko Takada

**Affiliations:** Systems Neuroscience Section, Department of Neuroscience, Primate Research Institute, Kyoto University, Inuyama, Aichi 484-8506, Japan; Department of Functional Brain Imaging, National Institutes for Quantum and Radiological Science and Technology, Chiba 263-8555, Japan; Laboratory for Cognitive Neuroscience, Graduate School of Frontier Biosciences, Osaka University, 1-4 Yamadaoka, Suita, Osaka 565-0871, Japan; Center for Information and Neural Networks, National Institute of Information and Communications Technology and Osaka University, 1-4 Yamadaoka, Suita, Osaka 565-0871, Japan; PRESTO, Japan Science and Technology Agency, Kawaguchi, Saitama 332-0012, Japan

**Author notes:** Co-corresponding authors: Ken-ichi Inoue, Masahiko Takada.

## Abstract

Recent emphasis has been placed on gene transduction mediated through recombinant adeno-associated virus (AAV) vector to manipulate activity of neurons and their circuitry in the primate brain. In the present study, we created a novel AAV vector of which capsid was composed of capsid proteins derived from the serotypes 1 and 2 (AAV1 and AAV2). Following the injection into the frontal cortex of macaque monkeys, this mosaic vector, termed AAV2.1 vector, was found to exhibit the excellence in transgene expression (for the AAV1 vector) and neuron specificity (for the AAV2 vector) simultaneously. To explore its applicability to chemogenetic manipulation and *in vivo* calcium imaging, the AAV2.1 vector expressing excitatory DREADDs or GCaMP was injected into the striatum or the visual cortex of macaque monkeys, respectively. Our results have defined that such vectors secure intense and stable expression of the target proteins and yield conspicuous modulation and imaging of neuronal activity.

## Introduction

In recent years, much attention has been attracted to manipulating activity of neurons and their circuitry by gene transduction with viral vectors (Adamantidis et al., 2007; Gradinaru et al., 2009; Zhang et al., 2010). Recombinant adeno-associated virus (AAV) vector has particularly been utilized for a variety of cutting-edge techniques, such as optogenetics and chemogenetics (for reviews, see Roth et al., 2016; Galvan et al., 2017; Atasoy et al., 2018). Indeed, substantial advances have been made in AAV vector-mediated gene transfer into target neuronal populations, and the validity of this strategy has been revealed in the primate brain as well as in the rodent brain (Alexander et al., 2009; Cavanaugh et al., 2012; Gerits et al., 2012; Jazayeri et al., 2012; Tye et al., 2012; Ohayon et al., 2013; Nagai et al., 2016). It is generally accepted that the AAV has numbers of serotypes, and that individual serotypes show different infectious properties (Zincarelli et al., 2008; Srivastava et al., 2016). In gene transfer experiments on the primate brain, vectors based on the serotypes 1, 5, and 9 of AAV (AAV1, AAV5, and AAV9) have so far frequently been used. These neurotropic AAV vectors exhibit higher levels of transgene expression than other serotype vectors, and, therefore, they may have been expected to exert adequate effects on modulating neuronal activity and animal’s behavior (Jazayeri et al., 2012; Klein et al., 2016; Stauffer et al., 2016; El-Shamayleh et al., 2017; Tamura et al., 2017). However, given that such vectors possess considerable infectivity to glial cells as well as to neurons, there seems to be a serious pitfall that they cause inflammatory responses due to the glial infection (Hadaczek et al., 2009; Markakis et al., 2010; Samaranch et al., 2014; Watakabe et al., 2015).

On the other hand, the serotype 2 of AAV (AAV2) is well known to display extremely high neuron specificity and, hence, low cytotoxicity (Fiandaca et al., 2008; Mueller et al., 2008; Russell et al., 2017; Domenger et al., 2019). Currently, recombinant vectors based on AAV2 are widely available for clinical use. Our previous works have demonstrated in macaque monkeys that optogenetic/chemogenetic manipulations mediated through the AAV2 vector successfully modulated oculomotor/cognitive behaviors of the animals as well as related neuron/pathway activities (Inoue et al., 2015; Nagai et al., 2016). However, these reports are rather exceptional on account of a lower transgene expression capacity of the AAV2 vector compared with other serotype vectors (Markakis et al., 2010; Watakabe et al., 2015). Thus, various serotypes of AAV vectors are being utilized in world-wide laboratories for gene transfer experiments, especially on the primate brain, and many neuroscientists are still seeking for an AAV vector optimal for manipulating target neuron/pathway activity. Conceivably, there is usually a trade-off relationship between the transgene expression capacity (i.e., sensitivity) and the neuron-specific infectivity (i.e., specificity). For improving the quality of gene transduction into the primate brain for diverse purposes, it should be essential to overcome this issue, thereby creating a novel AAV vector with marked superiority in both aspects.

Here, we developed an AAV vector of which capsid was composed of capsid proteins derived from both AAV1 and AAV2. This so-called mosaic vector, termed AAV2.1 vector, was designed to exhibit the excellence in transgene expression (for the AAV1 vector) and neuron specificity (for the AAV2 vector) simultaneously. By comparing with those of the original AAV2 and AAV1 vectors, we analyzed the gene transduction property of the AAV2.1 vector following the injection into the frontal cortex of macaque monkeys. In the present study, not only the transgene expression pattern of the mosaic vector, but also its applicability to chemogenetic manipulation and *in vivo* calcium imaging was explored in the striatum and visual cortex, respectively.

## Results

### Production of mosaic AAV vectors

For the present series of experiments, we produced two types of mosaic AAV vectors by changing the ratio of AAV1 and AAV2 capsid proteins as follows: AAV2.1-A vector with a combined capsid of 10% AAV1 capsid protein and 90% AAV2 capsid protein, and AAV2.1-B vector with a combined capsid of 50% AAV1 capsid protein and 50% AAV2 capsid protein. According to previous *in vitro* studies (Hauck et al., 2003; Rabinowitz et al., 2004; Choi et al., 2005), we initially made the AAV2.1-B vector. In our preliminary survey, however, we found that a certain degree of glial infectivity remained in this vector. Therefore, we subsequently made the AAV2.1-A vector in which the ratio of AAV1 capsid protein was decreased to a large extent (10%). A total of 14 different kinds of vectors based on the AAV2, AAV1, and their mosaic vectors was produced for three sets of our experiments. The production efficiency (stock titer) of each vector was as listed in Table 1. In general, AAV2 vectors had a lower efficiency than the others (i.e., AAV2.1 and AAV1 vectors).

**Table 1.** Summary of recombinant AAV vectors

| Animal ID | Serotype | % Capsid |  | Promoter | Gene | Stock titer<br>(gc/ml) | Injection<br>titer<br>(gc/ml) |
| --- | --- | --- | --- | --- | --- | --- | --- |
|  |  | AAV1 | AAV2 |  |  |  |  |
| Monkey A | AAV2 | 0 | 100 | CMV | mKO1 | $5.0 \times 10^{13}$ | $1.5 \times 10^{13}$ |
| Monkey B | | | | Syn | mKO1 | $8.0 \times 10^{13}$ | $6.0 \times 10^{13}$ |
| | AAV2.1-A | 10 | 90 | CMV | mKO1 | $2.2 \times 10^{14}$ | $1.5 \times 10^{13}$ |
| | | | | Syn | mKO1 | $2.8 \times 10^{14}$ | $6.0 \times 10^{13}$ |
| | AAV2.1-B | 50 | 50 | CMV | mKO1 | $9.3 \times 10^{14}$ | $1.5 \times 10^{13}$ |
| | | | | Syn | mKO1 | $1.0 \times 10^{15}$ | $6.0 \times 10^{13}$ |
| | AAV1 | 100 | 0 | CMV | mKO1 | $7.7 \times 10^{14}$ | $1.5 \times 10^{13}$ |
| | | | | Syn | mKO1 | $8.4 \times 10^{14}$ | $6.0 \times 10^{13}$ |
| Monkey C | AAV2 | 0 | 100 | Syn | hM3Dq-IRES-AcGFP | $1.8 \times 10^{13}$ | $1.0 \times 10^{13}$ |
| | AAV2 | 0 | 100 | CMV | hM3Dq-IRES-AcGFP | $1.0 \times 10^{13}$ | $1.0 \times 10^{13}$ |
| | AAV2.1-A | 10 | 90 | Syn | hM3Dq-IRES-AcGFP | $5.0 \times 10^{13}$ | $1.0 \times 10^{13}$<br>$5.0 \times 10^{13}$ |
| Monkey D | AAV2 | 0 | 100 | Syn | hM3Dq | $1.0 \times 10^{14}$ | $2.0 \times 10^{13}$ |
| | AAV2 | 0 | 100 | Syn | hM3Dq-IRES-AcGFP | $2.0 \times 10^{13}$ | $2.0 \times 10^{13}$ |
| | AAV2.1-A | 10 | 90 | Syn | hM3Dq-IRES-AcGFP | $4.0 \times 10^{13}$ | $2.0 \times 10^{13}$ |
| | AAV1 | 100 | 0 | Syn | hM3Dq-IRES-AcGFP | $8.9 \times 10^{13}$ | $2.0 \times 10^{13}$ |
| Monkey E | AAV2.1-A | 10 | 90 | CaMKIIα | GCaMP6s | $1.2 \times 10^{14}$ | $1.0 \times 10^{14}$ |
| | | | | | | | $2.0 \times 10^{13}$ |
| Monkey F | AAV2.1-A | 10 | 90 | CaMKIIα | GCaMP6s | $1.2 \times 10^{14}$ | $4.0 \times 10^{13}$ |

### Transgene expression patterns of mosaic AAV vectors

In the first series of our experiments, we investigated the gene transduction properties of the mosaic AAV2.1-A and AAV2.1-B vectors by comparing those of the original AAV2 and AAV1 vectors. To examine not only the transgene expression patterns of the mosaic vectors, but also the possible influences of promoters on their transgene expression, two types of promoters, ubiquitous cytomegalovirus (CMV) promoter and neuron-specific synapsin (Syn) promoter, were selected for evaluating the neuron specificity of the vectors and analyzing their transgene expression levels within neurons, respectively. To this end, injections of the following eight distinct kinds of AAV vectors carrying the mKO1 gene were made into the medial wall of the frontal lobe in two macaque monkeys (Monkeys A and B in Table 1): the AAV-CMV-mKO1 vectors decorated with four different compositions of capsids (AAV2, AAV2.1-A, AAV2.1-B, and AAV1) and the AAV-Syn-mKO1 vectors with the same virions (Fig. 1A, Table 1). All vectors loaded with CMV promoter were injected at the titer of 1.5×10^13^ genome copies (gc)/ml, while those loaded with Syn promoter were at the titer of 6.0×10^13^ gc/ml to make the transgene expression for the AAV2 vector detectable. We initially compared the intensity of transgene expression among these vectors at their cortical injection sites (Fig. 1B). The transgene expression level was quantified as an average value of the red fluorescent protein (RFP) signal within a circle of 2.0-mm diameter and depicted as the relative value to the expression level for the AAV2-CMV vector which was defined as 100 (Fig. 1C). With respect to the vectors loaded with CMV promoter, both the AAV1 and the AAV2.1-B vectors exhibited somewhat higher levels of transgene expression than the AAV2.1-A vector. Concerning the vectors loaded with Syn promoter, on the other hand, the AAV2.1-A vector displayed the highest expression level among the four vectors. Regardless of the promoter type, the transgene expression levels for the AAV2 vectors were so low as compared to the others. As for the CMV or Syn promoter, the expression level was only less than half or one-fifth as low as that for the AAV2.1-A vector, respectively (Fig. 1C).

**Fig. 1.**
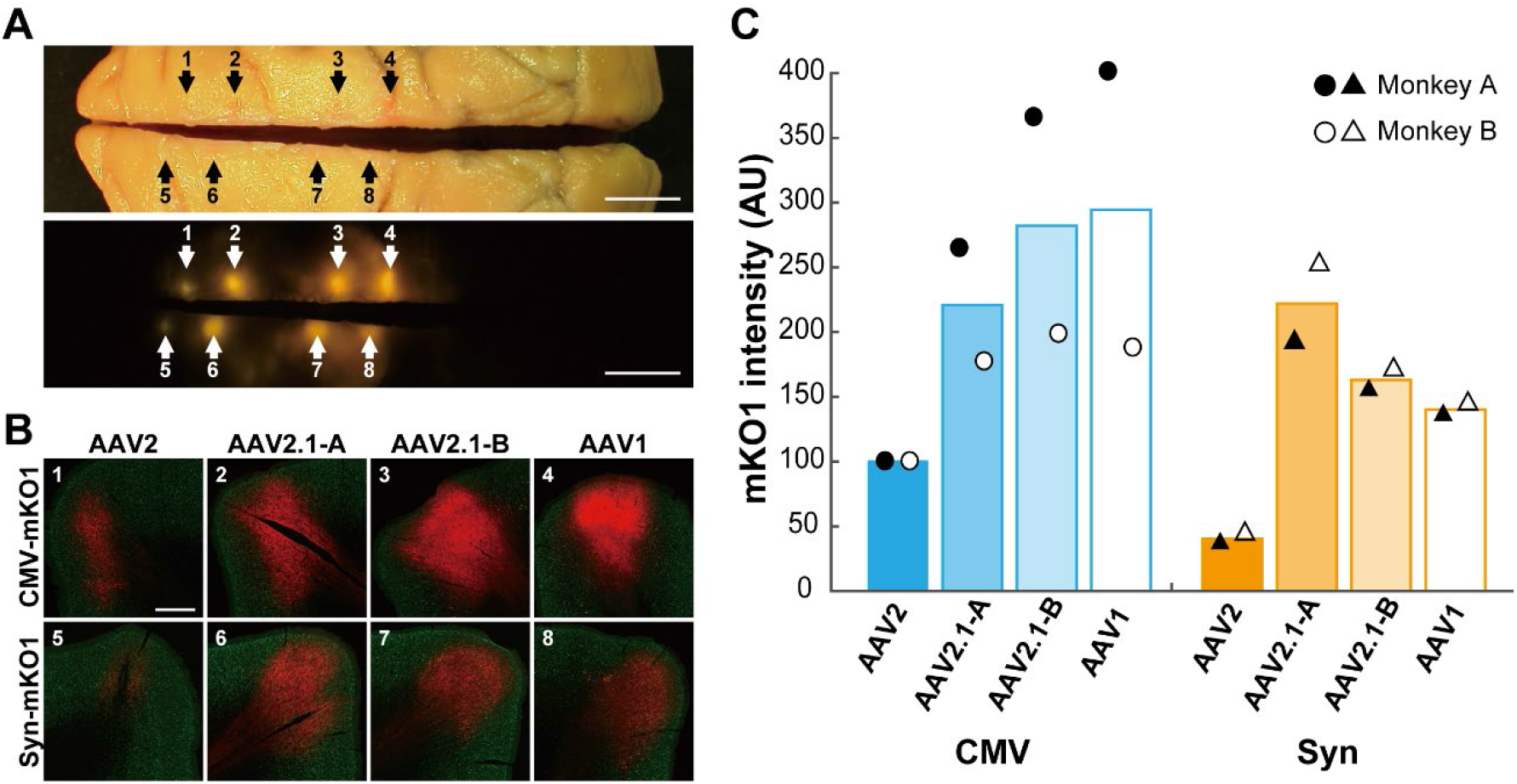
Intensity of mKO1 native-fluorescence at vector injection sites. (A) Injection loci of eight distinct types of recombinant AAV vectors carrying the mKO1 gene in the medial wall of the frontal lobe in Monkey A. Upper and lower rows represent bright-field image or mKO1 native-fluorescence image, respectively. Each arrow points to the injection site as follows: arrows 1-4, AAV2, AAV2.1A, AAV2.1B, and AAV1 vectors loaded with CMV promoter, respectively; arrows 5-8, AAV2, AAV2.1A, AAV2.1B, and AAV1 vectors loaded with Syn promoter, respectively. Scale bars, 10 mm. (B) mKO1 native-fluorescence (red) and NeuN immunofluorescence (green) at each vector injection site. Scale bar, 1 mm. (C) Intensity comparison of mKO1 native-fluorescence within a 2-mm diameter circle. Expressed as the mean value of data obtained in Monkeys A and B (denoted by filled and open circles and triangles), relative to the mean value for the AAV2-CMV vector defined as 1 arbitrary unit (AU).

Subsequently, we compared the mosaic vectors with the original vectors in terms of neuron specificity. Double fluorescence histochemistry was carried out to analyze the colocalization of NeuN as a neuronal marker or glial fibrillary acidic protein (GFAP) as an astroglial marker in RFP-positive (transduced) cortical cells (Fig. 2A,B). Cell counts were done by means of the stereological method, and the ratio of double-labeled cells to the total RFP-positive cells was calculated. Data on the vectors with CMV promoter showed that the neuron specificity of the AAV2.1-A vector was high enough to reach the level of the AAV2 vector, while the AAV2.1-B vector had certain glial infectivity, the extent of which was almost the same as the AAV1 vector (Fig. 2C). The vectors with Syn promoter exhibited almost complete or sufficiently high levels of neuron specificity, except that RFP expression in GFAP-positive (glial) cells remained for the AAV1 vector (Fig. 2C).

**Fig. 2.**
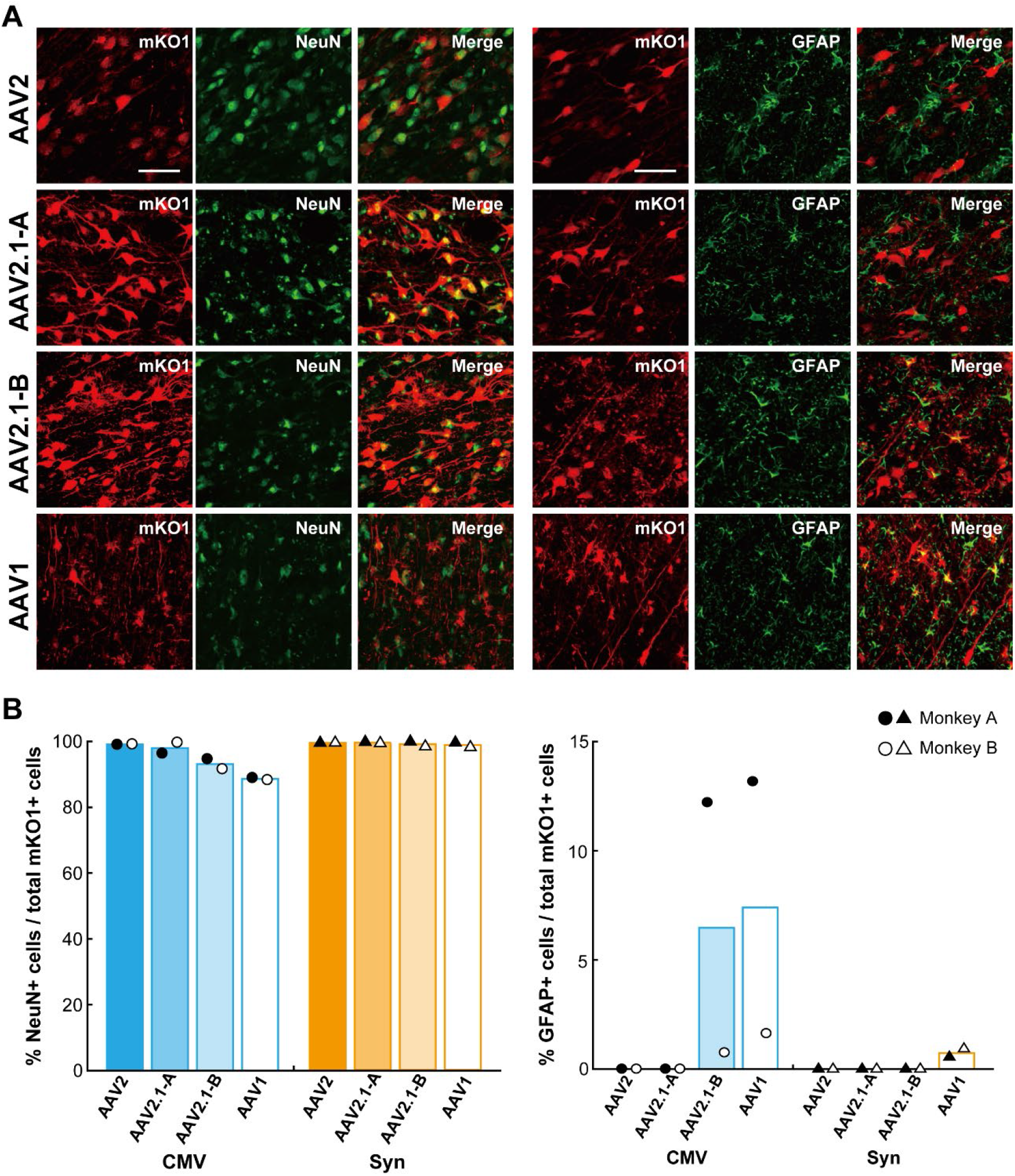
Transgene expression in neuronal and glial cells at vector injection sites. (A) Double fluorescence histochemistry for mKO1 native-fluorescence (red) and immunofluorescence for NeuN (green; left) or GFAP (green; right) at the injection sites of AAV vectors loaded with CMV promoter. Data in Monkey A. Shown in yellow (Merge) are double-labeled cells. Scale bars, 50 μm. (B) Ratio of double-labeled cells to the total mKO1-fluorescent cells at each vector injection site.

Overall, the AAV2.1-A vector resulted in the most efficient transgene expression with excellent neuron specificity, especially under the control of Syn promoter. It should also be noted that given their glial infectivity, the transgene expression images of the AAV1 and AAV2.1-B vectors with CMV promoter, which were more intense than that of the AAV2.1-A vector, might have involved the RFP signal issued not only from neuronal, but also from glial cells.

### Application of AAV2.1-A vector to chemogenetic manipulation

In the second series of our experiments, we explored the usefulness of the AAV2.1-A-Syn vector in chemogenetic manipulation of neuronal activity in the primate brain, by applying a recombinant vector carrying the hM3Dq gene to the excitatory DREADDs (designer receptors exclusively activated by designer drugs) system (Armbruster et al., 2007). The IRES-GFP (green fluorescent protein) sequence was further loaded as a fluorescent protein tag to examine the property of transduced neurons (AAV2.1-A-Syn-hM3Dq-IRES-GFP; see Table 1). For comparison, we also prepared four different vectors carrying the hM3Dq gene which were based on the original AAV2 or AAV1 vector and loaded with Syn or CMV promoter (AAV2-Syn-hM3Dq-IRES-GFP, AAV2-CMV-hM3Dq-IRES-GFP, AAV2-Syn-hM3Dq, and AAV1-Syn-hM3Dq-IRES-GFP; see Table 1). To test the potential of receptor binding, the responsiveness to ligand administration, and the duration of transgene expression, these vectors expressing the hM3Dq protein were injected into the striatum, the caudate nucleus and the putamen, of both hemispheres in two macaque monkeys (Table 1; see also Figs. 3A and 5A).

**Fig. 3.**
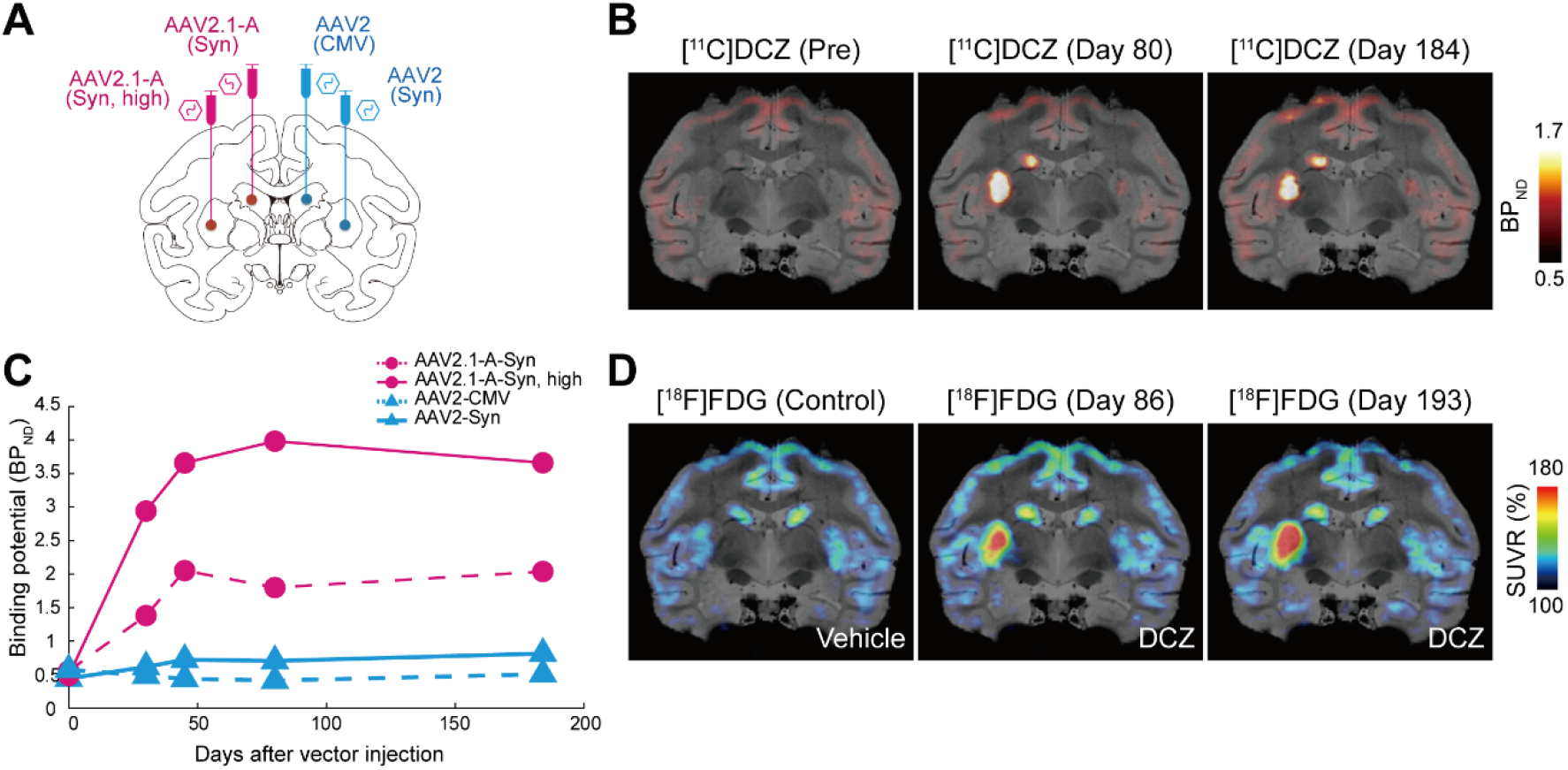
Chemogenetic manipulation of striatal neuron activity. (A) Sites of vector injections in the striatum in Monkey C. Four distinct vectors were injected into the striatum as follows: AAV2.1-A vector loaded with Syn promoter (Syn) into the caudate nucleus and the same vector at a higher titer (Syn, high) into the putamen on one side, and AAV2 vector loaded with CMV promoter (CMV) into the caudate nucleus and AAV2 vector loaded with Syn promoter (Syn) into the putamen on the other side. (B) Representative PET images overlaid on MR images showing [^11^C]DCZ-specific binding taken before (Pre) or at 80 and 184 days (Day 80, Day 184) after the vector injections. Each value of binding potential is expressed as the regional binding potential relative to a non-displaceable radioligand in tissue (BP_ND_). (C) Time-dependent changes of [^11^C]DCZ-specific binding at the sites of vector injections. Values before the injections, corresponding to (Pre) in (B), are plotted at Day 0. (D) Representative PET&MR-fused images showing normalized [^18^F]FDG uptake following vehicle administration (Control) or DCZ administration at 86 and 193 days (Day 86, Day 193) after the vector injections. Each value of FDG uptake is expressed as the standardized uptake value ratio (SUVR) to the mean value of the whole brain, averaged between 60 and 90 min after the radioligand injection.

In the first monkey (Monkey C in Table 1), to compare the AAV2.1-A vector with the AAV2 vector in terms of the three subjects listed above, we injected three types of highly neuron-specific vectors into the striatum as follows: the AAV2.1-A-Syn vector into the caudate nucleus and the putamen on one side, and the AAV2-CMV vector into the caudate nucleus and the AAV2-Syn vector into the putamen on the other side (Fig. 3A, Table 1). The injection titer of these vectors was set at 1.0×10^13^ gc/ml, except that only the AAV2.1-A-Syn vector was additionally injected at 5.0×10^13^ gc/ml into the putamen since its production efficiency (stock titer) was higher than those of the AAV2-CMV/Syn vectors (see Table 1). To evaluate the potential of DREADDs receptor binding *in vivo*, relevant to the strength of hM3Dq expression, positron emission tomography (PET) imaging with radiolabeled deschloroclozapine ([^11^C]DCZ) was carried out (see Nagai et al., 2020). To monitor the time-dependent changes in hM3Dq expression after intrastriatal injections of the AAV vectors, we performed [^11^C]DCZ-PET scans in the monkey at four times (30, 45, 80, and 184 days) post-injection (Fig. 3B,C). To verify the responsiveness to DREADDs ligand administration, we next attempted chemogenetic activation of neurons where hM3Dq was expressed through the AAV vectors. PET imaging with radiolabeled fluorodeoxyglucose ([^18^F]FDG) was carried out for *in vivo* visualization of glucose metabolism as an index of neuronal/synaptic activation (Phelps et al., 1979; Poremba et al., 2004; Michaelides et al., 2013). To detect the FDG uptake following systemic administration of DCZ, we performed [^18^F]FDG-PET scans in the monkey at twice (86 and 193 days) after the vector injections (Fig. 3D; see Nagai et al., 2020).

When the vectors were injected at the titer of 1.0×10^13^ gc/ml, the AAV2.1-A vector exhibited a sufficiently high level of [^11^C]DCZ binding (hM3Dq expression), whereas the [^11^C]DCZ binding levels for the AAV2 vectors were quite low regardless of the promoter type (Fig. 3B,C). In addition, the AAV2.1-A vector injected at a higher titer (5.0×10^13^ gc/ml) displayed an even greater level of [^11^C]DCZ binding compared with the lower-titer one (Fig. 3B,C). The [^11^C]DCZ-PET scans showed that the extent of [^11^C]DCZ binding for the AAV2.1-A vector was gradually increased until 45 days post-injection and then appeared to reach the plateau. Such hM3Dq expression lasted over six months (Fig. 3B,C). Similar results were obtained from the [^18^F]FDG-PET scans. At the injection site of the AAV2.1-A vector at the lower as well as the higher titer, the extent of FDG uptake was markedly increased by DCZ administration at 86 days after the vector injection and retained even at 193 days post-injection (Fig. 3D). On the other hand, virtually no increase was found for either type of the AAV2 vectors (Fig. 3D).

Subsequently, we histologically confirmed the PET imaging data described above. Immunohistochemical analysis with GFP antibody revealed that the localization of hM3Dq expression visualized by [^11^C]DCZ-PET imaging was in register with GFP-positive zones in the striatum, and that the GFP-positive zones corresponding to the injection sites of the AAV2.1-A vector were much denser or broader than those corresponding to the injection sites of the AAV2 vectors (Fig. 4A). Our GFP immunohistochemical analysis further showed that axon terminal labeling was evident in the substantia nigra (i.e., pars reticulata), whereas only very rarely was seen cell body labeling in the substantia nigra (i.e., pars compacta), thereby indicating that this vector was not preferentially transported in the retrograde direction (Fig. 4B). To verify neuronal activation chemogenetically induced by DCZ administration 2 hr before sacrifice, the density of striatal neurons expressing c-fos was examined at the injection site of each vector (Fig. 4C,D). Consistent with the findings obtained from [^18^F]FDG-PET imaging, the striatal neurons were activated by DCZ administration prominently at the injection site of the AAV2.1-A vectors including the higher-titer one, as compared to the AAV2 vectors.

**Fig. 4.**
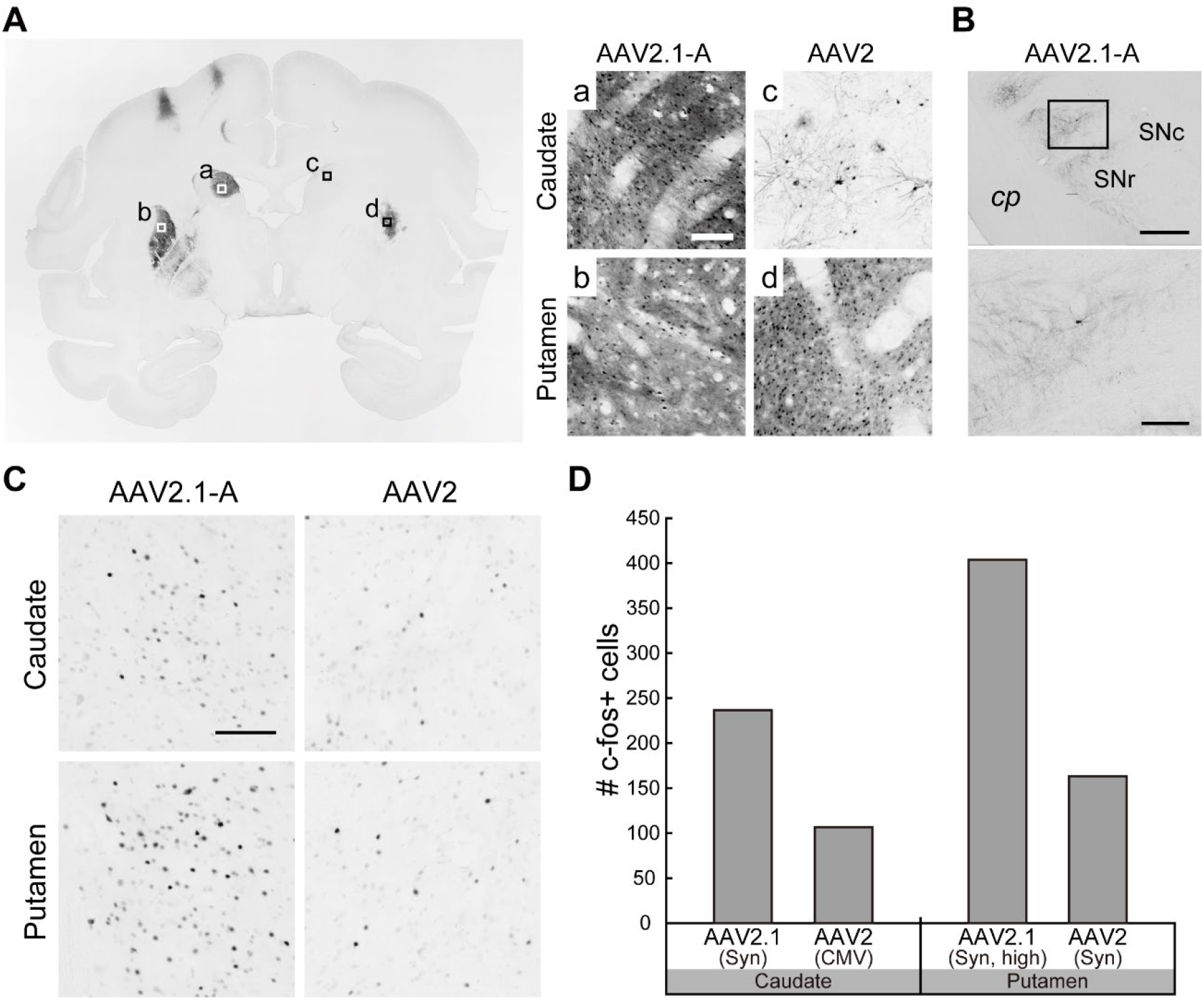
Histological analyses of striatal sections subjected to chemogenetic manipulation. (A) Left: Coronal section through the sites of vector injections immunostained with GFP antibody in Monkey C. Right: Higher-power magnifications of square areas a-d in the left panels. Scale bar, 150 μm. (B) GFP immunostaining showing terminal labeling in the substantia nigra on the side ipsilateral to the AAV2.1-A vector injection into the putamen. cp, cerebral peduncle; SNc, substantia nigra pars compacta; SNr, substantia nigra pars reticulata. Upper row: Scale bar, 1 mm. Lower row: Scale bar, 250 μm. (C) C-fos immunostaining at the injection sites of the vectors. Scale bar, 50 µm. (D) Number of c-fos-positive cells within a 1-mm diameter circle. Expressed as the mean value of cell counts obtained from three equidistant sections.

In the second monkey (Monkey D in Table 1), we injected four types of vectors including the AAV2.1-A vector into the striatum as follows: the AAV2-Syn vector into the caudate nucleus and the AAV2.1-A-Syn vector into the putamen on one side, and the AAV2-Syn vector without GFP tag into the caudate nucleus and the AAV1-Syn vector into the putamen on the opposite side (Fig. 5A, Table 1). The injection titer of these vectors was set at 2.0×10^13^ gc/ml. Since the AAV2 vector without fluorescent protein tag and the AAV1 vector have been used in previous DREADDs studies (Nagai et al., 2016, 2020), the validity of the AAV2.1-A vector was analyzed in comparison with these conventional vectors. In this monkey, [^11^C]DCZ-PET scans were carried out at seven times (30, 45, 64, 120, 143, 176, and 358 days) after the vector injections. Similar high levels of [^11^C]DCZ binding (hM3Dq expression) were observed for all of the AAV2 vector without GFP tag, the AAV1 vector, and the AAV2.1-A vector (Fig. 5B,C). The binding levels were gradually increased until 64 days post-injection and then reached the plateau. Such hM3Dq expression was retained as long as one year (Fig. 5B,C). With respect to the FDG uptake, we performed [^18^F]FDG-PET scans at five times (49, 72, 129, 182, and 365 days) after the vector injections and observed that the AAV2.1-A vector, together with the AAV2 vector without GFP tag and the AAV1 vector, displayed a sufficiently high level of FDG uptake induced by DCZ administration (Fig. 5D). Such responsiveness of striatal neurons to the DREADDs ligand also lasted over one year. Apparently, the extent of FDG uptake for the AAV2 vector without GFP tag seemed even higher than the AAV2.1-A and AAV1 vectors. Consistent with the findings obtained in the first monkey, the AAV2-Syn vector with GFP tag exhibited much lower levels of [^11^C]DCZ binding and FDG uptake than the other three vectors (Fig. 5D).

### Application of AAV2.1-A vector to in vivo calcium imaging

In the third series of our experiments, we investigated the applicability of the AAV2.1-A vector to *in vivo* calcium imaging in the primate brain. A recombinant vector carrying the GCaMP6s gene which was loaded with CaMKIIα promoter (AAV2.1-A-CaMKIIα-GCaMP6s; see Table 1) was produced and injected into the primary and secondary visual cortical areas (V1 and V2) of two macaque monkeys. Since the first monkey (Monkey E in Table 1) was used primarily for determining optimal experimental conditions, data obtained in the second monkey (Monkey F in Table 1) as a representative were shown below. We initially performed intrinsic signal optical imaging to visualize an orientation map of the visual cortex, in which response signals to visual stimuli on various directions were monitored under general anesthesia. Then, both one-photon wide-field calcium imaging and two-photon calcium imaging were carried out to capture signals of neuronal activity from distinct areas (Fig. 5A). Time-dependent changes in fluorescent signals were recorded at 41 and 244 days after the vector injection (Fig. 5B). The fluorescent signals obtained from many neurons were robust at both sampling days. Even at 244 days post-injection, the captured signals did not appear to be attenuated in comparison with those at 41 days post-injection. In addition, explicit orientation-selective responses of V1 and V2 neurons to drifting grating stimuli could be recorded at both 41 and 244 days post-injection (Fig. 5C). These overall results indicated that high levels of transgene expression in the visual cortex via the AAV2.1-A vector were maintained over eight months.

**Fig. 5.**
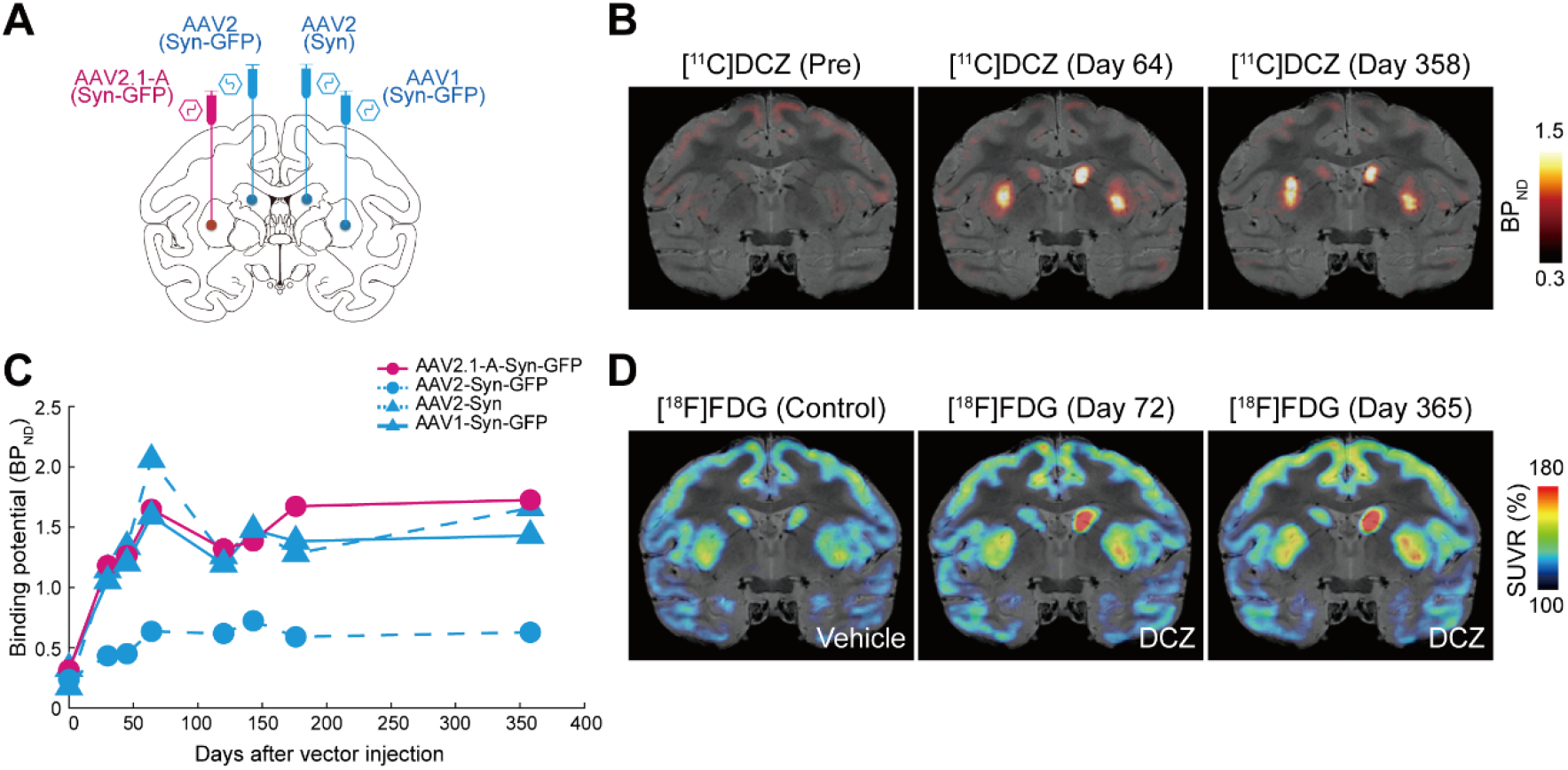
Chemogenetic manipulation of striatal neuron activity. (A) Sites of vector injections in the striatum in Monkey D. Four distinct vectors were injected into the striatum as follows: AAV2 vector with Syn promoter and GFP tag (Syn-GFP) into the caudate nucleus and AAV2.1-A vector with Syn promoter and GFP tag (Syn-GFP) into the putamen on one side, and AAV2 vector with Syn promoter but without GFP tag (Syn) into the caudate nucleus and AAV1 vector with Syn promoter and GFP tag (Syn-GFP) into the putamen on the other side. (B) Representative PET images overlaid on MR images showing [^11^C]DCZ-specific binding (expressed as BP_ND_) taken before (Pre) or at 64 and 358 days (Day 64, Day 358) after the vector injections. (C) Time-dependent changes of [^11^C]DCZ-specific binding at the sites of vector injections. Values before the injections, corresponding to (Pre) in (B), are plotted at Day 0. (D) Representative PET&MR-fused images showing normalized [^18^F]FDG uptake (expressed as SUVR) following vehicle administration (Control) or DCZ administration at 72 and 365 days (Day 72, Day 365) after the vector injections.

**Fig. 6.**
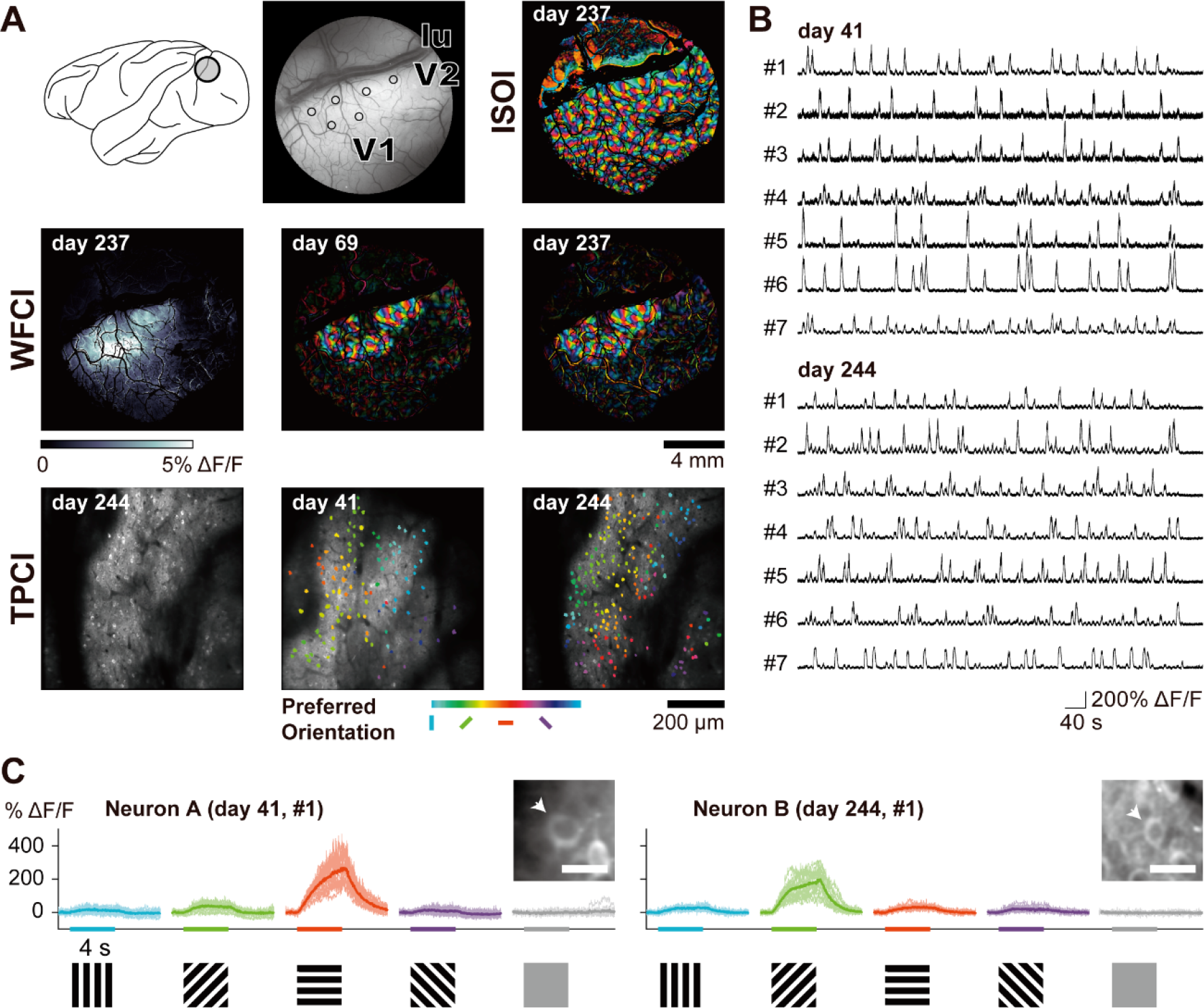
*In vivo* calcium imaging of visual cortical neuron activity. (A) Upper row: Left, a recording chamber placed over the V1 and V2 in Monkey F; Central, six loci of AAV2.1-A vector (AAV2.1-A-CaMKIIα-GCaMP6s) injections (denoted by open circles). lu, lunate sulcus; Right, intrinsic signal optical imaging (ISOI) for visualizing an orientation map of the visual cortex at 237 days post-injection. Middle row: One-photon wide-field calcium imaging (WFCI). Left, vector injection sites at 237 days post-injection visualized by selecting the maximum fluorescence changes (ΔF/F) for each pixel across all stimulus conditions; Central & Right, orientation maps visualized at 69 and 237 days post-injection. The color code indicates the preferred orientation determined for each pixel. Note that these maps are basically consistent with the one obtained from ISOI (see the upper right) and, also, at the two timepoints apart over six months. Lower row: Two-photon calcium imaging (TPCI). Left, structural image of the recording sites at 244 days post-injection obtained by averaging the fluorescence across all frames for a recording session. Central & Right, orientation maps at the single-neuron resolution at 41 and 244 days post-injection. The color code represents the preferred orientation determined for each neuron. (B) Time-dependent changes (ΔF/F) in fluorescent signals obtained from 14 exemplified neurons in the V1 recorded at 41 and 244 days after the vector injection. (C) Responses of fluorescent signals of two V1 neurons (Neurons A and B) to drifting gratings of different orientations. The responses to the same orientation with opposite moving directions are superimposed. (Insets) Arrowheads point to the imaged neurons with typical nucleus-empty appearance of GCaMP labeling. Scale bars, 20 μm.

## Discussion

In the present study, we have newly developed the mosaic AAV vector, termed AAV2.1 vector, of which capsid was composed of capsid proteins derived from both AAV1 and AAV2. Of the two types of mosaic vectors with different compositions of their capsid proteins, the AAV2.1-A vector of which capsid was obtained from a higher ratio of AAV2 capsid protein (10% AAV1 and 90% AAV2) has successfully met the two requirements, high levels of transgene expression and neuron specificity. Our detailed analysis has revealed that this vector possesses the excellence in transgene expression (for the AAV1 vector) and neuron specificity (for the AAV2 vector) simultaneously. Concerning the vectors loaded with CMV promoter, the transgene expression level of the AAV2.1-A vector was somewhat lower than those of the AAV1 vector and the AAV2.1-B vector (another mosaic vector with a capsid obtained from the same ratio of AAV1 and AAV2 capsid proteins). When the CMV promoter was replaced with the Syn promoter, however, the AAV2.1-A vector exhibited the highest level of transgene expression. Based on the fact that the AAV1 and AAV2.1-B vectors with CMV promoter had glial infectivity to a certain extent, their transgene expression images might have involved the fluorescent signals issued not only from neuronal, but also from glial cells. Thus, it is most likely that transgene expression through many recombinant AAV-CMV vectors may represent signals issued from both neuronal and glial cells, although these two signals cannot technically be discriminated. On the other hand, the expression levels of the AAV1 and AAV2.1-B vectors with Syn promoter were so limited as compared to the AAV2.1-A vector since neither the AAV1 nor the AAV2.1-B vector was able to express the transgene under the control of Syn promoter, even though infection to glial cells occurred.

To explore its applicability to chemogenetic manipulation and calcium imaging in the primate brain, the AAV2.1-A vector expressing excitatory DREADDs or GCaMP was injected into the striatum or the visual cortex, respectively. Our results have verified that the AAV2.1-A vector secures intense and stable expression of the target protein and achieves unequivocal modulation and imaging of neuronal activity. In the present experiments on chemogenetic manipulation, we initially compared the AAV2.1-A vector with the AAV2 vector. The overall data clearly demonstrated that the AAV2.1-A vector displayed sufficiently high levels of both the potential of DREADDs receptor binding and the responsiveness to DREADDs ligand administration. It should also be noted that the AAV2.1-A vector employed at a higher titer yielded a greater capacity of transgene expression. The productivity of such a high-titer vector might be ascribable to an AAV1 component contained in this vector, because the production efficiency of AAV1 vector is relatively high to that of AAV2 vector. On the other hand, the transgene expression capacity of the AAV2 vector was still lower regardless of the promoter type even compared with the AAV2.1-A vector at the same titer. In striking contrast, the capacity of the AAV2 vector without florescent protein tag became much higher, especially in terms of the responsiveness to DREADDs ligand administration. It is generally accepted that insertion of the IRES-GFP sequence into conventional AAV vectors dampens transgene expression (Zhou et al., 1998; Mizuguchi et al., 2000; Fuler et al., 2001). However, given its potentially high level of transgene expression (see “Transgene expression patterns of mosaic AAV vectors” in the Results section), the AAV2.1-A vector can be considered to retain a sufficient expression level even with the florescent protein tag loaded. On any occasion, AAV vectors without florescent protein tag do not appear to merit application to many studies which require anatomical confirmation of vector localization/transport or histological determination of the type/number of transduced cells.

By comparing the AAV2.1-A vector with the AAV1 vector, we subsequently found that the transgene expression level of the AAV1 vector was equivalent to that of the AAV2.1-A vector. This is not necessarily surprising, because AAV1 vector has been regarded as a stronger transgene carrier for the primate brain than AAV2 vector (Watakabe et al., 2015). For the AAV1 vector to secure neuron-specific gene expression, however, the selection of a promoter, like Syn promoter, is critical. Therefore, the usefulness of this vector would be rather restricted depending on the promoter type.

For applying viral vectors to gene transfer experiments on the primate brain, their sufficiently high levels of transgene expression are indispensable. In this context, multiple vectors such as AAV1, AAV5, and AAV9 vectors have often been utilized (Jazayeri et al., 2012; Klein et al., 2016; Stauffer et al., 2016; El-Shamayleh et al., 2017; Tamura et al., 2017). Nevertheless, since these vectors cause inflammatory responses due to their glial transduction (Hadaczek et al., 2009; Markakis et al., 2010; Gray et al., 2011; Ciesielska et al., 2013; Samaranch et al., 2014; Watakabe et al., 2015; Tanabe et al., 2019; Shirley et al., 2020; Verdera et al., 2020; present results), presumable tissue damage following intracranial injections of the vectors largely discourages their application to long-term experimental studies over years in primates. The present work has successfully overcome this issue by creating the AAV2.1-A vector, a novel mosaic vector that allows to thoroughly diminish glial infectivity and potentially reduce inflammatory responses, thereby permitting behavioral and electrophysiological monitoring over a long period of time after neuron/pathway activity manipulation induced by target gene transduction.

For recent years, many neuroscientists have made an effort to gain reliable and stable modulation of neuronal activity and animal behavior in primates by combining the viral vector system with various cutting-edge techniques. The AAV2.1-A vector developed in our study bears a prominent advantage over the conventional AAV vectors in view of the fact that this vector is endowed with the excellence in both transgene expression and neuron specificity in the primate brain, and that both the AAV2 and the AAV1 vectors likely have potential limitations on their utility and applicability, as stated above. Such a mosaic AAV vector can be useful as a versatile tool for diverse *in vivo* approaches, including chemogenetic manipulation and calcium imaging we tested here. The AAV2.1-A vector exhibits the excellent neurotropism even without Syn promoter. This enables us to replace the Syn promoter with other types of promoters so as to modify the transgene expression pattern or achieve the neuron type-selective gene transduction. The widespread and effective application of the AAV2.1-A vector not only could facilitate elucidating neural network mechanisms underlying higher-order brain functions, but also might subserve establishing valid primate models of neurological/psychiatric disorders.

## Subject details

### Animals

Six adult macaque monkeys were used for the present study: one rhesus monkey (*Macaca mulatta*), 5.6 kg, female; one cynomolgus monkey (*Macaca fascicularis*), 5.2 kg, male; four Japanese monkeys (*Macaca fuscata*), 5.3-8.3 kg, one male, three females (Monkeys A-F in Table 1). For the first series of our experiments in Monkeys A and B (i.e., Transgene expression patterns of mosaic AAV vectors), the experimental protocol was approved by the Animal Welfare and Animal Care Committee of the Primate Research Institute, Kyoto University (Permission Number: 2018-046), and all experiments were conducted according to the Guidelines for Care and Use of Nonhuman Primates established by the Primate Research Institute, Kyoto University (2010). For the second series of our experiments in Monkeys C and D (i.e., Application of AAV2.1-A vector to chemogenetic manipulation), the experimental protocol was approved by the Animal Ethics Committee of the National Institutes for Quantum and Radiological Science and Technology (Permission Number: 11-1038-11), and all experiments were conducted in accordance with the Guide for the Care and Use of Laboratory Animals (National Research Council of the US National Academy of Sciences). For the third series of our experiments in Monkeys E and F (i.e., Application of AAV2.1-A vector to in vivo calcium imaging), the experimental protocol was approved by the Animal Experiment Committee of Osaka University (Permission Number: FBS-18-005), and all experiments were conducted according to the Guidelines for Animal Experiments established by Osaka University.

## Method details

### Viral vector production

All AAV vectors were produced by the helper-free triple transfection procedure. Briefly, AAV-293 cells (70% confluent in Corning Cell Stack 10 chamber) were transfected by genome, helper (pHelper; Stratagene, SanDiego, USA), and packaging plasmids (pAAV-RC1 and/or pAAV-RC2) with polyethylenimine (PEI Max; Polysciences, Warrington, USA). For production of AAV2.1-A or AAV2.1-B vector, the pAAV-RC1 plasmid-coding AAV1 capsid protein and the pAAV-RC2 plasmid-coding AAV2 capsid protein were transfected respectively with the ratio of 1:9 or 5:5 (Hauck et al., 2003; Rabinowitz et al., 2004; Choi et al., 2005). The produced vectors were then purified by affinity chromatography (GE Healthcare, Chicago, USA), concentrated to 150 μl by ultrafiltration (Amicon Ultra-4 10K MWCO; Millipore, Billerica, USA), and the titer of each vector stock was determined by quantitative PCR using Taq-Man technology (Life Technologies, Waltham, USA). The transfer plasmid (pAAV-CMV-mKO1-WPRE) was constructed by inserting the mKO1 gene (MBL Lifescience, Tokyo, Japan) and the WPRE sequence into an AAV backbone plasmid (pAAV-CMV; Stratagene).

The pAAV-hSyn-mKO1-WPRE plasmid was constructed by replacing the CMV promoter with the human Syn promoter, and, further, the pAAV-Syn-hM3Dq-WPRE plasmid was constructed by replacing the mKO1 gene with the hM3Dq gene. Construction of the pAAV-Syn-hM3Dq-IRES-AcGFP-WPRE plasmid was as reported elsewhere (Nagai et al., 2016, 2020), and the pAAV-CMV-hM4Di-IRES-AcGFP-WPRE plasmid was constructed by promoter replacement as described above. The pAAV-CaMKIIα-GCaMP6s-WPRE plasmid was constructed by replacing the CMV promoter and the mKO1 gene of the pAAV-CMV-mKO1-WPRE plasmid with the mouse CaMKIIα promoter (0.4 k base pair) and the GCaMP6s gene, respectively.

### Cortical injections of viral vectors

Two monkeys (Monkeys A and B in Table 1) were initially sedated with ketamine hydrochloride (5 mg/kg, i.m.) and xylazine hydrochloride (0.5 mg/kg, i.m.), and then anesthetized with sodium pentobarbital (20 mg/kg, i.v.) or isoflurane (1-3%, to effect). An antibiotic (Ceftazidime; 25 mg/kg, i.v.) was administered at the initial anesthesia. After removal of a skull portion over the frontal lobe, eight different types of AAV vectors expressing RFP were injected bilaterally into the medial wall of the frontal lobe by the aid of a magnetic resonance imaging (MRI)-guided navigation system (Brainsight Primate; Rogue Research, Montreal, Canada). The injections were made through a 10-μl Hamilton microsyringe (0.5 μl/site, two sites per track, one track for each vector). After the experiments, the monkeys were monitored until the full recovery from the anesthesia. All experimental procedures were performed in a special laboratory (biosafety level 2) designated for *in vivo* animal infectious experiments that had been installed at the Primate Research Institute, Kyoto University. Throughout the entire experiment, the animals were kept in individual cages that were placed inside a special safety cabinet in controlled temperature (23-26°C) and light (12-hr on/off cycle) conditions. The animals were fed regularly with dietary pellets and had free access to water. Every effort was made to minimize animal suffering.

### Chemogenetic manipulation

Under general anesthesia as described above, five distinct types of AAV vectors carrying the hM3Dq gene were injected bilaterally into the striatum (the caudate nucleus and the putamen) in two other monkeys (Monkeys C and D in Table 1). Stereotaxic coordinates of the injected sites were defined based on overlaid magnetic resonance (MR) and computed tomography (CT) images created by PMOD image analysis software (PMOD Technologies, Zurich, Switzerland). The intrastriatal injections were made in a similar manner to that described above (3 μl/site, one site per track, one track for each vector). Afterward, PET imaging was performed using the same procedures as described previously (Nagai et al., 2020). Briefly, the monkeys were sedated with ketamine hydrochloride (5 mg/kg, i.m.) and xylazine hydrochloride (0.5 mg/kg, i.m.), and the anesthetized condition was maintained with isoflurane (1-2%) during the PET imaging. PET scans were done with microPET Focus220 scanner (Siemens Medical Solutions, Malvern, USA). Following transmission scans, emission scans were acquired for 90 min after intravenous bolus injection of [^11^C]DCZ (331.5-404.9 MBq) or [^18^F]FDG (233.3-306.9 MBq). Pretreatment with DCZ (1 μg/kg; MedChemExpress, Monmouth Junction, USA) or vehicle (1-2% dimethyl sulfoxide (DMSO) in 0.1-µl saline, without DCZ) was carried out 1 min before the [^18^F]FDG injection. The PET imaging data were reconstructed with filtered back-projection with attenuation correction. Voxel values were converted to standardized uptake values (SUVs) that were normalized by injected radioactivity and body weight using PMOD. Volumes of interest (VOIs) were manually drawn on the center of the injection site and the cerebellum using PMOD, referring to MR images of individual monkeys. To estimate the specific binding of [^11^C]DCZ, the regional binding potential relative to nondisplaceable radioligand (BP_ND_) was calculated with an original multilinear reference tissue model using the cerebellum as a reference region (Ichise et al., 1996). For FDG–PET analysis, dynamic SUV images were motion-corrected and then averaged between 60 and 90 min after the radioligand injection. The SUV ratio (SUVR) of voxel value was calculated as a percentage to the mean value of the whole brain for comparison across the scans.

### In vivo calcium imaging

In the two remaining monkeys (Monkeys E and F in Table 1), we performed *in vivo* intrinsic signal optical imaging (ISOI), wide-field calcium imaging (WFCI) and two-photon calcium imaging (TPCI) of neurons in layers 2 and 3 of the V1 and V2. Under sterile conditions and isoflurane anesthesia (1-2%), the cortical surface of the V1 and V2 was exposed by incision of the scalp and removal of a skull portion and dura over these areas. The AAV2.1-A vector carrying the GCaMP6s gene was injected unilaterally into the V1 and V2 at six sites (1 μl each site). At each site, two injections were made at the depth of 1 and 2 mm (0.5 µl each depth). After the vector injections, a chronic titanium imaging chamber (inner diameter, 12 mm) was implanted to cap the exposed cortex. The center of the chamber was positioned 15-mm lateral to the midline, and its anteroposterior position was adjusted so that the lunate sulcus ran along the one-third upper position of the chamber to allow a view of both the V1 and the V2 (Fang et al., 2018). The exposed cortex was covered with artificial dura (W. L. Gore & Associates, Inc., Newark, USA) that was glued to the chamber with silicone adhesive (Kwik-Sil; World Precision Instruments, Sarasota, USA) for preventing regenerative tissue from infiltrating into the imaging field (Li et.al., 2017). The exposed cortex was further protected with a cover glass (Matsunami, Osaka, Japan; 14 mm in diameter and 0.5 mm in thickness), which was fixed to the chamber with a retaining ring (SM14RR; Thorlabs, Newton, USA). The retaining ring was glued to the titanium chamber with surgical silicone adhesive (Kwik-Sil).

When we performed imaging experiments, anesthesia was first introduced to the monkeys with ketamine and then maintained with propofol (induction, 5 mg/kg; maintenance, 5 mg/kg/hr, i.v.; Fang et al., 2018). The monkeys were paralyzed with vecuronium bromide (induction, 0.25 mg/kg, i.v.; maintenance, 0.05 mg/kg/hr, i.v.) and artificially respirated. The eyes were fitted with contact lenses of appropriate curvature to prevent from drying and focus on a stimulus screen 57-cm apart from the cornea (Ikezoe et al., 2018). In ISOI, intrinsic signals were acquired by a CMOS camera tandem-lens setup (FLIR GS3-U3-41C6NIR-C; exposure time, 100-300 μs; aperture, f5.6; lens combination, Noct-Nikkor 58 mm for the object side and Nikkor 50 mm for the CMOS side; Nikon, Tokyo, Japan) with 625-nm illumination (M625L3; Thorlabs) at a frame rate of 25 frames/s. One single imaging trial lasted 5 s (1 s before visual stimulus onset and 4 s after onset). The image size was 1024 × 1024 pixels representing a 12.5 ×12.5 mm^2^ field of view, which covered the full field of the cortical surface within the imaging chamber. Images were acquired with custom-built software made with LABVIEW (National Instruments, Austin, USA). The setup for WFCI was similar to that for ISOI and shared the same imaging window and software. Calcium signals were acquired by the same CMOS camera (exposure time, 10-30 ms) with 490-nm illumination (M490L4; Thorlabs) and camera-side 500-nm filtering (FELH0500; Thorlabs) at a frame rate of 25 frames/s and lasted 5 s. In TPCI, calcium signals were acquired with a two-photon microscope (MOM with 8 kHz resonant scanner; Sutter, Novato, USA) equipped with a 16×, NA 0.8 water immersion objective lens (CFI175-LWD-16XW; Nikon) and 920-nm excitation laser (Insight X3; Spectra Physics, Milpitas, USA). The frame rate was 30.9 frames/s. The size of image areas was 512 × 512 pixels representing a 630 × 630 μm^2^ field of view.

Visual stimuli were created using ViSaGe Mk II (Cambridge Research Systems, Rochester, UK) and displayed on a gamma-calibrated 23-inch LCD monitor (MDT231WG; Mitsubishi Electronic, Tokyo, Japan). We presented full-screen drifting black-white square-wave gratings. The duty cycle of gratings was 0.2 (20% white), and the spatial frequency was 1.5 cycle/degree. The gratings drifted at 8°/s and were presented in a randomly interleaved fashion in four orientations (0°, 45°, 90°, and 135°) with two opposite moving directions (i.e., eight directions; 0°, 45°, 90°, 135°, 180°, 225°, 270°, and 315°). For a blank screen condition, there were nine stimulus conditions in total. Each stimulus condition lasted 4 s on screen. The initial phase of the gratings was randomly selected.

In data analysis, fluorescent response (ΔF/F) was first quantified using the following formula: ΔF_i_/F = (F_i_ - F_0_)/F_0_, in which F_i_ was a response in an *i*-th single frame after stimulus onset, and F_0_ was an average response of frames over a period of 1 s before stimulus onset. We used this fluorescent response to describe the response time course of a population of single neurons. In ISOI and WFCI, we compared responses to orthogonal orientations by performing subtraction of the averaged ΔF/F between two orthogonal orientation conditions. For this analysis, responses to opposite directions with the same orientation gratings were averaged. All image processing procedures were done with the MATLAB (Mathworks). Blood vessel mask was calculated based on the cortical image under 530-nm illumination (M530L3; Thorlabs).

### Histology and image acquisition

Monkeys A-D underwent perfusion-fixation. The approximate survival periods after the vector injections were four weeks for Monkeys A and B, over six months for Monkey C, and over 12 months for Monkey D. In Monkey C, DCZ was administered systemically 2 hr prior to sacrifice for analysis of c-fos expression. The animals were anesthetized deeply with an overdose of sodium pentobarbital (50 mg/kg, i.v.) and perfused transcardially with 0.1 M phosphate-buffered saline (PBS; pH 7.4), followed by 10% formalin in 0.1 M phosphate buffer. The removed brains were postfixed in the same fresh fixative overnight at 4°C and saturated with 30% sucrose in 0.1 M PBS at 4°C. Coronal sections were cut serially at the 50-µm thickness on a freezing microtome and divided into ten groups. Every tenth section was used for individual histological analyses.

Double fluorescence histochemistry for RFP and NeuN or GFAP was performed as described in our prior study (Tanabe et al., 2019). Briefly, coronal sections were incubated with mouse monoclonal antibody against NeuN (1:2,000; Sigma-Aldrich, St. Louis, USA) or GFAP (1:1,000; Sigma-Aldrich). These sections were then incubated with Alexa 647-conjugated donkey anti-mouse IgG antibody (1:400; Jackson ImmunoResearch, West Grove, USA). For immunohistochemical staining for GFP and c-fos following chemogenetic manipulation, coronal sections were pretreated with 0.3% H_2_O_2_ for 30 min, rinsed three times in 0.1 M PBS, and immersed in 1% skim milk for 1 hr. The sections were then incubated for two days at 4°C with rabbit monoclonal anti-GFP antibody (1:2,000; Invitrogen, Waltham, USA) or rabbit polyclonal anti-c-fos antibody (1:2,000; abcam, Cambridge, UK) in 0.1 M PBS containing 2% normal donkey serum and 0.1% Triton X-100. Subsequently, the sections were incubated with biotinylated donkey anti-rabbit IgG antibody (1:1,000; Jackson ImmunoResearch) in the same fresh medium for 2 hr at room temperature, followed by the avidin-biotin-peroxidase complex (ABC *Elite*; 1:200; Vector laboratories, Burlingame, USA) in 0.1 M PBS for 2 hr at room temperature. The sections were finally reacted for 10-20 min in 0.05 M Tris-HCl buffer (pH 7.6) containing 0.04% diaminobenzine tetrahydrochloride (Wako, Osaka, Japan), 0.04% NiCl_2_, and 0.002% H_2_O_2_. The reaction time was set to make the density of background immunostaining almost identical. These sections were mounted onto gelatin-coated glass slides and counterstained with 0.5% Neutral red.

A high sensitivity charge-coupled device (CCD) camera (VB-7010, Keyence, Osaka, Japan) was used to capture bright-field and fluorescent images from the whole brain sample. To capture the bright-field and fluorescent images in coronal sections, a scientific CMOS camera (In Cell Analyzer 2200; GE Healthcare, Chicago, USA) was used. We created merged images of mKO1 and NeuN/GFAP-immunofluorescent coronal sections by using Fiji/ImageJ software (Schindelin et al., 2012). A confocal laser-scanning microscope (LSM800; Carl Zeiss, Oberkochen, Germany) was also used to take fluorescent images and prepare merged images using the Zeiss Zen 3.3 software (Carl Zeiss).

## Quantification and statistical analysis

### Intensity analysis of mKO1 native-fluorescence at injection sites

The intensity of mKO1 native-fluorescence issued from each vector injection site was measured by using Matlab software (Mathworks, Natick, USA). Three sections were chosen including the sections through the center of the injection site and 500 µm anterior/posterior to the injection site. The brightest circle (2 mm in diameter) was defined as a region of interest (ROI) in each section, and the mean value of fluorescence intensity obtained from three ROIs was calculated. To estimate the transgene expression level at each injection site, the expression level for the AAV2-CMV vector was defined as 100, and the relative value to this was calculated for the other vectors.

### Transgene expression analysis in neuronal and glial cells at injection sites

Three sections were chosen in the same fashion as described above. Stereological cell counts assisted with Stereo Investigator software (MBF Biosciences, Williston, USA) were carried out in 100-μm×100-μm counting frames equally spaced across a 250-μm×250-μm grid were counted with an 18-μm-high optical dissector (average section thickness: 20-μm). Based on such data, the ratio of double-labeled cells to the total RFP-positive cells was calculated.

### Counts of c-fos-positive cells

Three sections were chosen in the same manner as described above. At each vector injection site in the striatum, the number of c-fos positive cells was counted by the aid of Neurolucida software (MBF Biosciences). The densest circle (1 mm in diameter) of c-fos positive cells was defined as a ROI in each section, and the mean value of cell counts obtained from three ROIs was calculated.

## Acknowledgements

We are grateful to K. Nagaya, E. Tanaka, E. Sumiya, R. Suma, Y. Sugii, R. Yamaguchi, Y. Matsuda, J. Kamei, and N. Nitta for their technical assistance. We also thank Dr. M.-R. Zhang and his colleagues at the Department of Radiopharmaceuticals Development, QST, for producing the radioligand. This work was supported by MEXT/JSPS KAKENHI Grant Numbers JP19F193830 to Y.F., JP14J01649 to R.T., JP18H05007 and JP21H02596 to I.F., JP19H03335 to K.I., and JP19H05467 to M.T., by AMED Grant Numbers JP21dm0107146 to T.M., JP20dm0307021 to K.I., and JP21dm0207077 to M.T., and by JST Grant Number JPMJCR1683 to K.I.

## Author contributions

K.I. and M.T. designed the experiments. M.F., M.N., and K.I. prepared the viral vectors. K.K., S.T., A.Z., and K.I. performed the experiments on transgene expression pattern analysis of mosaic vectors. K.K., Y.N., Y.H., and T.M. performed the experiments on the application of the mosaic vectors to chemogenetic manipulation. G.H., Y.F., R.T., M.I., I.F., and K.I. performed the experiments on the application of the mosaic vectors to *in vivo* calcium imaging. K.K., Y.N., and G.H. analyzed the data. K.K., T.M., I.F., K.I., and M.T. wrote the manuscript. All authors discussed the data and commented on the manuscript.

